# Dual roles of glycine betaine (GB), dimethylglycine, and sarcosine as osmoprotectants and nutrient sources for *Vibrio natriegens*

**DOI:** 10.1101/2025.03.18.643870

**Authors:** Heather E. Thomas, Katherine E. Boas Lichty, Gary P. Richards, E. Fidelma Boyd

## Abstract

Bacteria respond to osmotic stress by intracellularly accumulating low molecular weight compounds called compatible solutes (CS), also known as osmolytes. Glycine betaine (*N*,*N*,*N*-trimethylglycine, GB) is a highly effective and widely available osmolyte used by bacteria, algae, and plants for abiotic stress protection. Here, we highlight the dual roles of GB, dimethyl glycine (DMG), and sarcosine for both osmoprotection and a less known role as sole carbon sources. First, we showed that the marine halophile *Vibrio natriegens* can grow in 1% to 7% NaCl and biosynthesize GB, ectoine, and glutamate, and imported GB, DMG, and sarcosine in response to osmotic stress. Betaine-carnitine-choline transporters (BCCTs) for the uptake of GB and DMG, but not sarcosine, were identified. Bioinformatics analyses uncovered homologs of GB, DMG, and sarcosine catabolism genes (*dgcAB_fixAB, gbcA, gbcB, purU, soxBDAG, glyA, glxA*) clustered in the *V. natriegens* genome and these genes had a limited distribution among vibrios. We showed *V. natriegens* ATCC 14048 grew on GB, DMG, and sarcosine as sole carbon sources and *gbcA* and *dgcA* were required for growth. A contiguous catabolism cluster was present in a subset of *V. fluvialis* strains, and we demonstrated growth of *V. fluvialis* 2013V-1197 in DMG and sarcosine as sole carbon sources. Phylogenetic analysis revealed the catabolism cluster did not share a common ancestor among members of the family *Vibrionaceae*.

**IMPORTANCE:** Compatible solutes are frequently the most concentrated organic components in marine organisms allowing them to adapt to high saline environments as well as affording protection to other abiotic stresses. These organic compounds are significant energy stores that have been overlooked for their potential as abundant nutrient sources for bacteria. Our study characterized GB, DMG, and sarcosine catabolism genes and showed their efficient use as carbon and energy sources by marine halophilic vibrios.

## INTRODUCTION

Marine bacteria encounter a range of salt concentrations in their environment and must account for osmotic stress and the resulting changes in turgor pressure. To compensate for osmotic stress, bacteria have evolved different mechanisms in order to maintain cellular homeostasis (1–9). A common and widely used strategy is the intracellular accumulation of compatible solutes, also known as osmolytes (1, 2, 5, 7–10). Compatible solutes are small organic compounds that can be accumulated in molar concentrations without causing disruptions to essential cellular functions (1, 2, 5, 7–10). In hyper-osmotic environments, osmolytes are accumulated intracellularly by either uptake from the environment or biosynthesis. There are only a few examples of *de novo* osmolyte biosynthetic pathways present in bacteria, such as ectoine biosynthesis from its endogenous precursor L-aspartate. Most biosynthetic pathways require a precursor from the surrounding environment such as choline for glycine betaine (GB) biosynthesis (encoded by *betIBA*) (11–13). Transport of osmolytes into the cell is accomplished using specialized osmolyte transporters, members of the ATP-binding cassette (ABC) family that require ATP, secondary transporters of the Betaine Carnitine Choline Transporter (BCCT) family, the Major facilitator superfamily (MFS), and the tripartite ATP-independent periplasmic (TRAP) family of transporters (14–24). Osmolyte transporters are predominantly expressed under high NaCl conditions and repressed when the osmotic stress is removed (17, 20–22, 24–27).

Previous studies characterized osmolyte uptake by *Vibrio cholerae*, *V. coralliilyticus*, *V. fluvialis, V. harveyi*, *V. parahaemolyticus,* and *V. vulnificus,* showing all species utilized GB as a highly effective osmoprotectant and biosynthesized GB from choline (21, 25, 28–32). Studies in *V. parahaemolyticus* showed optimal growth in up to 6% NaCl, and the ability to biosynthesize ectoine, glutamate, and GB, and to uptake 15 different osmolytes under osmotic stress conditions, which included choline, (GB), *N*,*N*-dimethylglycine (DMG), dimethylsulfoniopropionate (DMSP), and ectoine using high affinity BCCT transporters for uptake (11, 21, 25, 29–31, 33).

Osmolytes are organic compounds and therefore have the potential to be used as carbon or nitrogen, and energy sources (13, 30, 34). GB is an abundant osmolyte produced and utilized by bacteria, plants, and animals for stress protection (35–41).

Levels of GB in marine environments range from picomolar in seawater, nanomolar on particulate matter to mM range in phytoplankton and 100 mM range in corals (34–41). While the ability of bacteria to biosynthesize GB from choline is ubiquitous, their capacity to catabolize GB is much less common (13, 30, 34). A foundational study examining soil and activated sludge samples demonstrated that a large number of aerobic bacteria could utilize choline, GB, DMG and sarcosine as sole carbon and nitrogen sources (42). That study showed that osmolyte degradation was found mainly in coryneforms and *Pseudomonas* genera (42). Aerobic catabolism of GB was subsequently demonstrated in *Corynebacterium*, *Arthrobacter*, *Sinorhizobium, Chromohalobacter, Halomonas*, and *Pseudomonas* species (17, 43–54). The aerobic catabolism pathway involved serial demethylation of GB to dimethylglycine (DMG) to methylglycine (sarcosine) to glycine (**Fig. 1**) (17, 50, 52, 55–57). The first gene in the pathway is *gbcA* which encodes a Rieske-family oxidase, while a related gene, *gbcB,* encodes a ferredoxin reductase recently described in *Chromohalobacter salexigens,* which together convert GB to DMG and formaldehyde (58, 59). In *Sinorhizobium meliloti,* this first step in GB catabolism involves the enzyme betaine homocysteine methyltransferase (BHMT), which converts GB and homocysteine to DMG and methionine (46, 47, 60). The next step is DMG conversion to sarcosine requiring dimethylglycine demethylase (DDM) encoded by *dgcAB,* which is followed by sarcosine conversion to glycine by a tetrameric sarcosine oxidase (TSOX) encoded by *soxBDAG* in *Pseudomonas* species (**Fig. 1**). Glycine is further broken down by serine hydroxymethyltransferase (*glyA1*) and serine dehydratase (*sda*) to pyruvate to enter central metabolism (**Fig. 1**). In *P. aeruginosa*, transcription of *gbcA, gbcB, dgcAB,* and *soxBDAG* requires the activators GbdR, an AraC transcription regulator, which is induced by GB and DMG, and SouR, which is induced by sarcosine (52, 56, 57, 61). Recent marine microbial community studies examining the fate of exogenous GB using metabolomics and transcriptomic analyses highlighted GB uptake and catabolism as an important source of carbon, nitrogen, and energy (62–65). Previous bioinformatic analyses identified homologs of several enzymes required for GB catabolism in *V. natriegens*, a marine halophile first isolated in the salt marsh mud of Sapelo Island, Georgia, USA and a member of the ubiquitous Harveyi clade (30, 66).

**Fig. 1.**
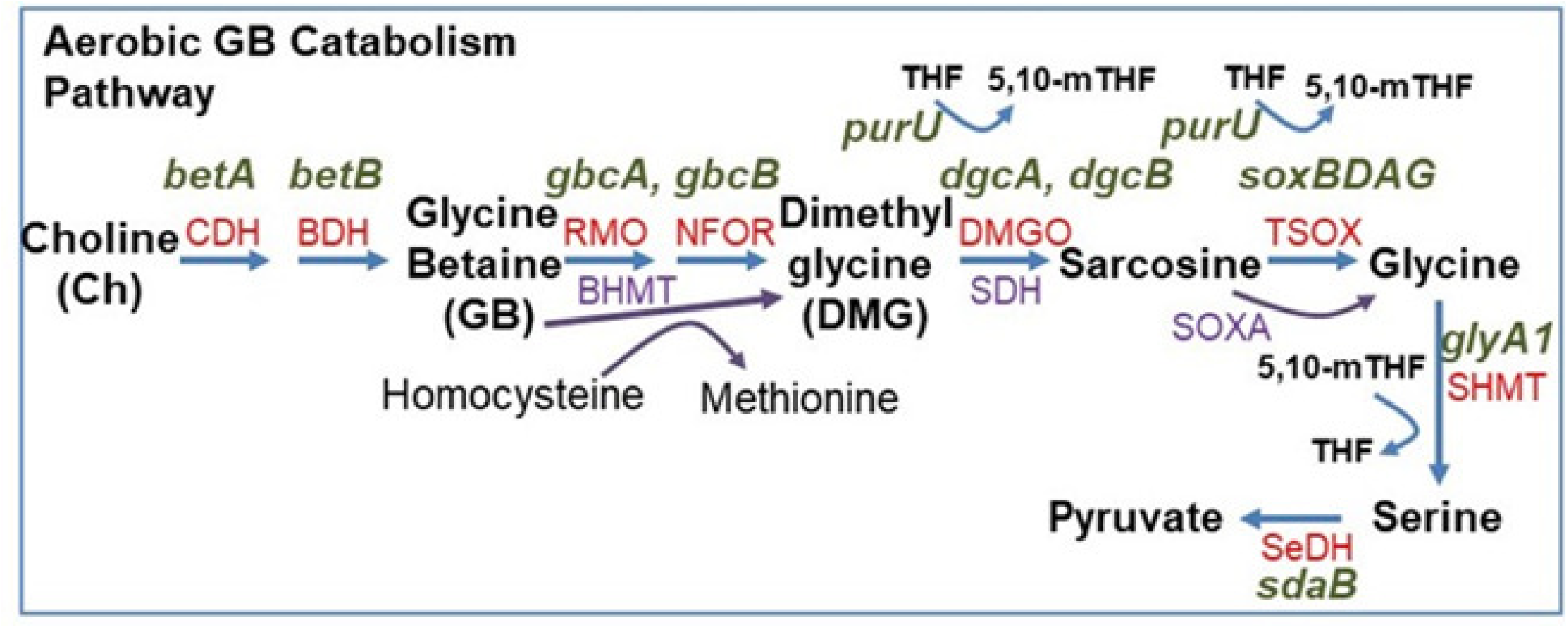
Glycine betaine aerobic catabolism pathway. Enzymes abbreviations in red and gene designations in black above are those present in *Pseudomonas aeruginosa*. Abbreviations: aerobic pathway in red, choline dehydrogenase (CDH, *betA*), betaine aldehyde dehydrogenase (BDH, *betB*), Rieske mono-oxygenase (RMO, *gbcA*), NADH:Flavin oxidoreductase (NFOR, *gbcB*), DMG oxidases (DMGO,*dgcAB*), heterotetrameric sarcosine oxidase (TSOX, *soxBDAG*). Glycine can be further broken down by serine hydroxymethyltransferase (SHMT, *glyA1*) and serine dehydratase (SeDH, *sdaB*) to pyruvate to enter central metabolism. Tetrahydrofolate (THF); 5,10-methylenetetrahydrofolate (5,10-mTHF). Enzyme abbreviations in purple represent alternative enzymes present in other bacteria such as *Sinorhizobium meliloti* uses betaine-homocysteine methyltransferase (BHMT) to convert GB and homocysteine into DMG and methionine. Some *Bacillus* species use monomeric sarcosine oxidase (SOXA) to convert sarcosine to glycine.

In this study, we describe the osmotic stress response of *V. natriegens* to determine its NaCl tolerance and the mechanisms it uses to adapt to salinity. Members of the family *Vibrionaceae* have not been shown previously to utilize GB, DMG, or sarcosine as carbon and energy sources, which was also examined here. First, we determined the range of salinities *V. natriegens* could grow in and whether NaCl played a role in temperature tolerance. Next, we examined the range of osmolytes that *V. natriegens* could utilize in response to osmotic stress. Using a bioinformatic approach, we examined the genome of *V. natriegens* ATCC 14048 for homologs of previously identified osmotic stress systems. Proton nuclear magnetic resonance (^1^H-NMR) spectroscopy, was used to determine the compatible solute biosynthesized by this species. Osmolyte uptake assays of stressed cells were used to examine osmolyte transport and the role individual BCCT transporters played. Genome context and gene neighborhood analyses identified a BCCT transporter clustered with genes showing homology to enzymes involved in the conversion of GB to pyruvate. The ability to consume GB, DMG and sarcosine as carbon and energy sources was examined in *V. natriegens* and *V. fluvialis*. The distribution of the catabolism cluster was examined among *Vibrionaceae*.

## RESULTS

### *Vibrio natriegens* response to salinity and temperature

*Vibrio natriegens* ATCC 14048 was grown in lysogeny broth (LB) 3% NaCl (wt./vol.) at 37°C for 24-h. A short lag phase was observed with the cultures growing to an optical density (OD_595_) of ∼1.55 in less than 2.5-h, with a growth rate of 1.9 confirming previous studies of it fast growth rate (data not shown) (66–68). To determine the NaCl tolerance range of *V. natriegens*, we examined growth in M9 minimal media supplemented with 20 mM glucose (M9G) and 0% NaCl to 7.5% NaCl in the absence of external osmolytes at 30°C. *Vibrio natriegens* grew optimally in 0% to 5% NaCl; whereas at 6% and 6.5% NaCl, they grew after extended lag phases, while at 7% NaCl, no growth was observed (**Fig. 2A**). At 37°C, *V. natriegens* grew optimally between 1% and 6% NaCl, at 7% NaCl it had an extended lag phase; and at 7.5% NaCl there was no growth (**Fig. 2B**). In addition, *V. natriegens* did not grow in the absence of NaCl at 37°C. These data indicate that *V. natriegens* has a higher NaCl tolerance range at 37°C, but also has a requirement for NaCl to grow at this temperature indicating that NaCl plays a role in thermal tolerance.

**Fig. 2.**
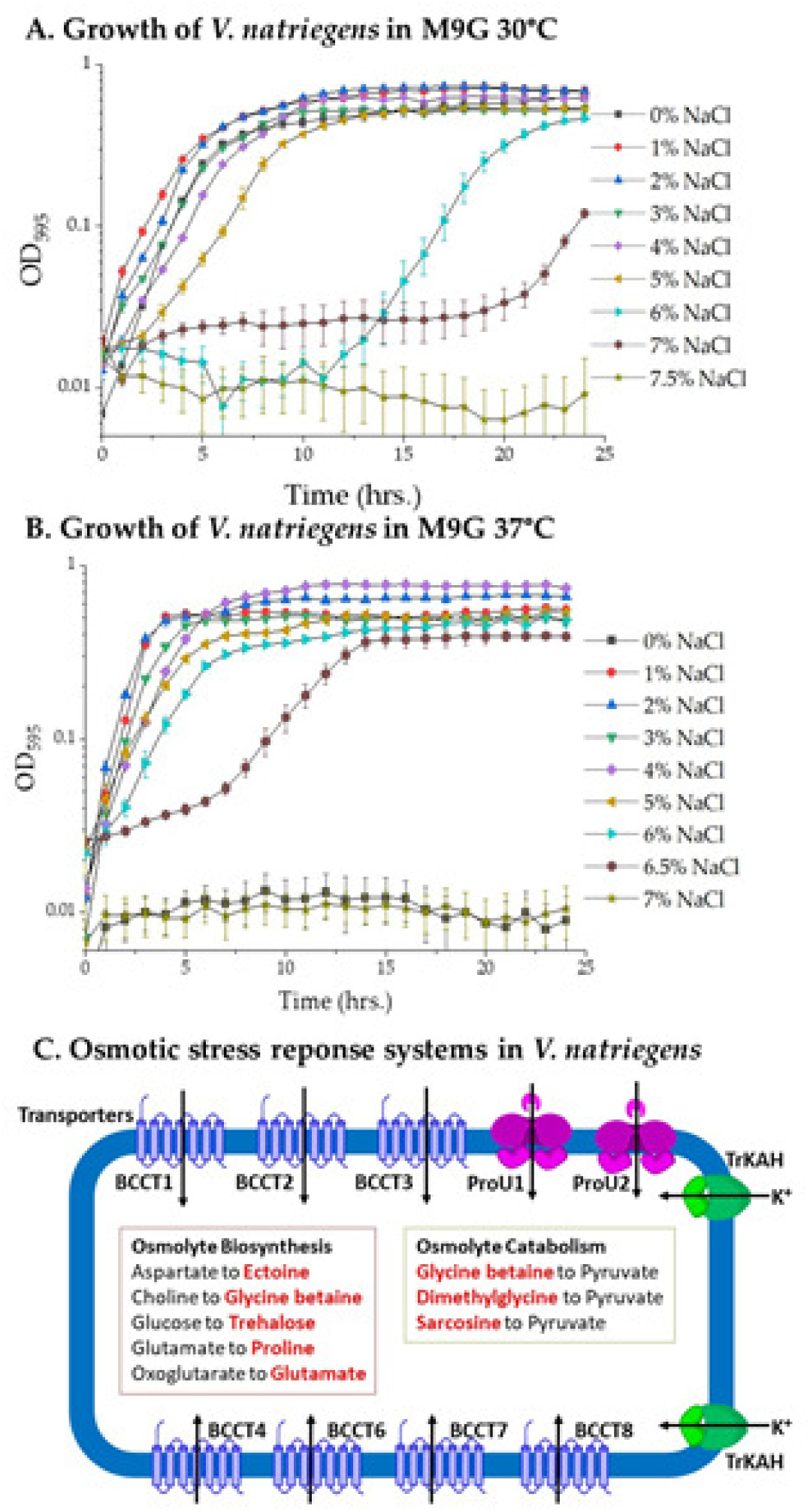
*Vibrio natriegens* ATCC 14048 NaCl tolerance range. Growth curves of *V. natriegens* ATCC 14048 in minimal media (M9) with glucose (M9G) with a final concentration of 1% to 7.5% NaCl. Growth was measured every hour for 24 h at **A.** 30°C, **B.** 37°C and **C**. Osmotic stress response systems identified in *V. natriegens* ATCC 14048. Depicted are two TrkAH transporters for the uptake of potassium a short-term response to osmotic stress, seven BCCT transporters identified, two ABC-type transporters, ProU1 and ProU2 encode by *proVWX.* Also shown are osmolyte biosynthesis pathways and osmolyte catabolic pathways identified. Arrows show the direction of substrate transport.

### Osmotic stress response systems in *V. natriegens*

Bioinformatics analysis was performed comparing osmotic stress systems present in *V. parahaemolyticus* RIMD 2210633 to the *V. natriegens* ATCC 14048 genome RefSeq GCF_001456255.1. BLAST analysis uncovered two putative osmolyte biosynthesis operons for ectoine (encoded by *ectABC_aspK*) and glycine betaine (GB) (encoded by *betIBA*) as well as nine putative osmolyte transporters (**Fig. 2C**). Chromosome I (NZ_CP009977.1) contained two betaine-carnitine-choline transporters (BCCTs), BCCT1 and BCCT3 and a ProU transporter (encoded by *proVWX)*, which was adjacent, but divergently transcribed from *ectABC_aspK*. Chromosome II (NZ_CP009978.1) contained five BCCTs, BCCT2 (PN96_RS23610), BCCT4 (PN96_RS17645), BCCT6 (PN96_RS15305), BCCT7 (PN96_RS21680), and BCCT8 (PN96_RS22655), and a ProU transporter (*proXWV*, PN96_RS19360-PN96_RS19370) in an operon *betIBA_proXWV*. The numbering of BCCT1 to BCCT4 is based on their homology to the BCCTs from *V. parahaemolyticus* RIMD 2210633. BCCT5 was identified previously in *Vibrio vulnificus* and is absent from *V. natriegens* (29). Therefore, when numbering the additional BCCTs present in *V. natriegens*, BCCT5 was skipped. Overall, the *V. natriegens* ATCC 14048 genome contains seven BCCTs whereas stains CL-2, TC2-11, and SY.PD55 contained eight BCCTs, four more than are present in *V. parahaemolyticus*.

### Ectoine, glutamate, and glycine betaine are biosynthesized by *V. natriegens* for osmotic stress protection

The putative ectoine biosynthesis operon *ectABC*_*aspK* should allow for the biosynthesis of ectoine *de novo* from endogenous aspartic acid (**Fig. 2C**). To exam this, proton nuclear magnetic resonance (^1^H-NMR) spectroscopy was performed on ethanol extracts of *V. natriegens* grown in M9G 5% NaCl. The spectral peaks corresponding to the various hydrogen atoms in the ectoine molecule are labelled in **Fig. 3A**. This analysis also identified peaks corresponding to the osmolyte glutamate, indicating it is also produced in response to osmotic stress (**Fig. 3A**). As a negative control, cells were also grown in M9G 1% NaCl, and extracts from these unstressed cells lacked ectoine and glutamate proton peaks (**Fig. S1**). Next, *V. natriegens* was examined for its ability to biosynthesize GB from its exogenously supplied precursor choline. ^1^H-NMR spectra of ethanol extracts of *V. natriegens* grown in M9G 5% NaCl supplemented with 1 mM choline showed signal peaks corresponding to GB (**Fig. 3B**). Ectoine peaks were absent in these spectra, demonstrating ectoine is not biosynthesized in the presence of choline (**Fig. 3B**). Overall, the data showed that *V. natriegens* was able to biosynthesize both glutamate and ectoine *de novo* and GB in the presence of choline in response to osmotic stress.

**Fig. 3.**
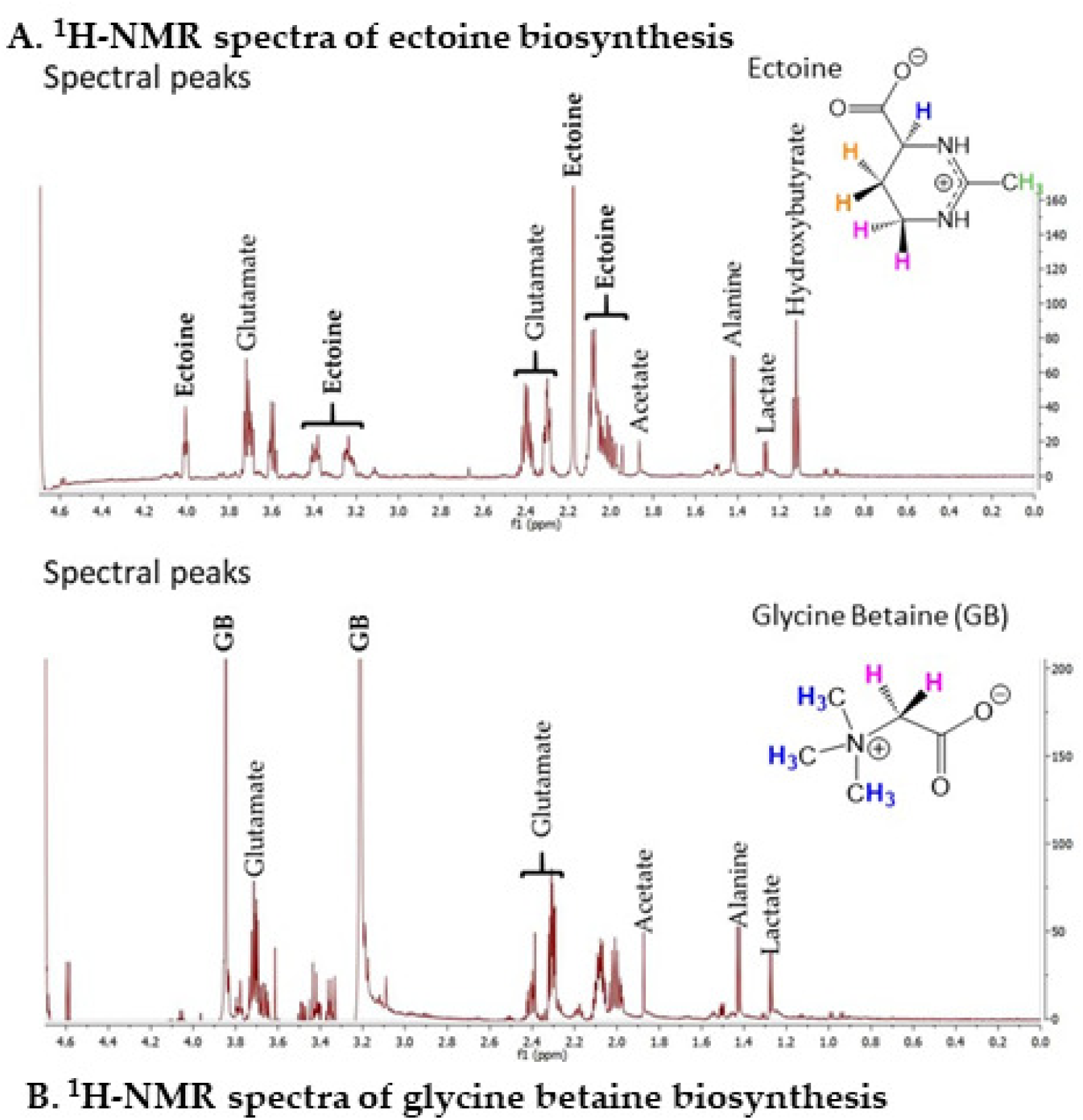
**Biosynthesis of compatible solutes in *Vibrio natriegens* ATCC 14048. A. ^1^**H-NMR spectra of *V. natriegens* ATCC 14048 cellular extract grown in minimal media (M9G) 5%NaCl and **B.** M9G 5%NaCl with the addition of 1 mM choline. The spectral peaks corresponding to glutamate, ectoine and glycine betaine (GB) are labeled. The protons spectral peaks corresponding to **A.** glutamate (peaks at 3.7 ppm, 2.4 ppm, 2.3 ppm) and ectoine (4.0 ppm, 3.4 ppm, 3.25 ppm, 2.1-2.2 ppm singlet, 2.0-2.1 ppm multilet) and **B.** GB (3.85 ppm, 3.2 ppm) and glutamate (peaks at 3.7 ppm, 2.4 ppm, 2.3 ppm) are labeled, as illustrated by their chemical shift values expressed in ppm on the x-axis.

### *Vibrio natriegens* can uptake and utilize a range of osmolytes for osmoprotection

To determine the range of osmolytes taken up and utilized by *V. natriegens* for osmoprotection, cells were grown in M9G 7% NaCl supplemented with 1 mM of choline, GB, DMG, sarcosine, DMSP, ectoine, or L-proline. In the absence of osmolytes, there was a 6-h lag phase, which was significantly reduced in the presence of 1 mM of all solutes examined, with GB, DMG, and DMSP showing the greatest lag phase reduction (**Fig. 4A**). Next, these osmolytes were examined to determine whether they restored growth in M9G 7.5% NaCl, a condition that inhibits growth. M9G 7.5% NaCl was supplemented with 100 μM of each osmolyte, and GB, DMG, DMSP, choline, and L-proline rescued growth. The length of the lag phase for each substrate suggests there may be differences in transporter affinities. For example, GB has the shortest lag phase indicating it is taken up efficiently whereas L-proline has a 5-h lag phase suggesting a less efficient transporter. Ectoine and sarcosine did not rescue growth at this concentration (**Fig. 4B**). When the concentration was increased to 1 mM, all seven osmolytes rescued growth with sarcosine having the longest lag phase, but rescues to wild-type levels (**Fig. 4C**). This suggests that sarcosine does not have an efficient transporter for uptake in high NaCl. Overall, these data indicate that *V. natriegens* was able to uptake and utilize osmolytes as osmoprotectants, and that choline, GB, DMG, and DMSP are most effective based on their shorter lag phases and ability to rescue growth at low concentrations.

**Fig. 4.**
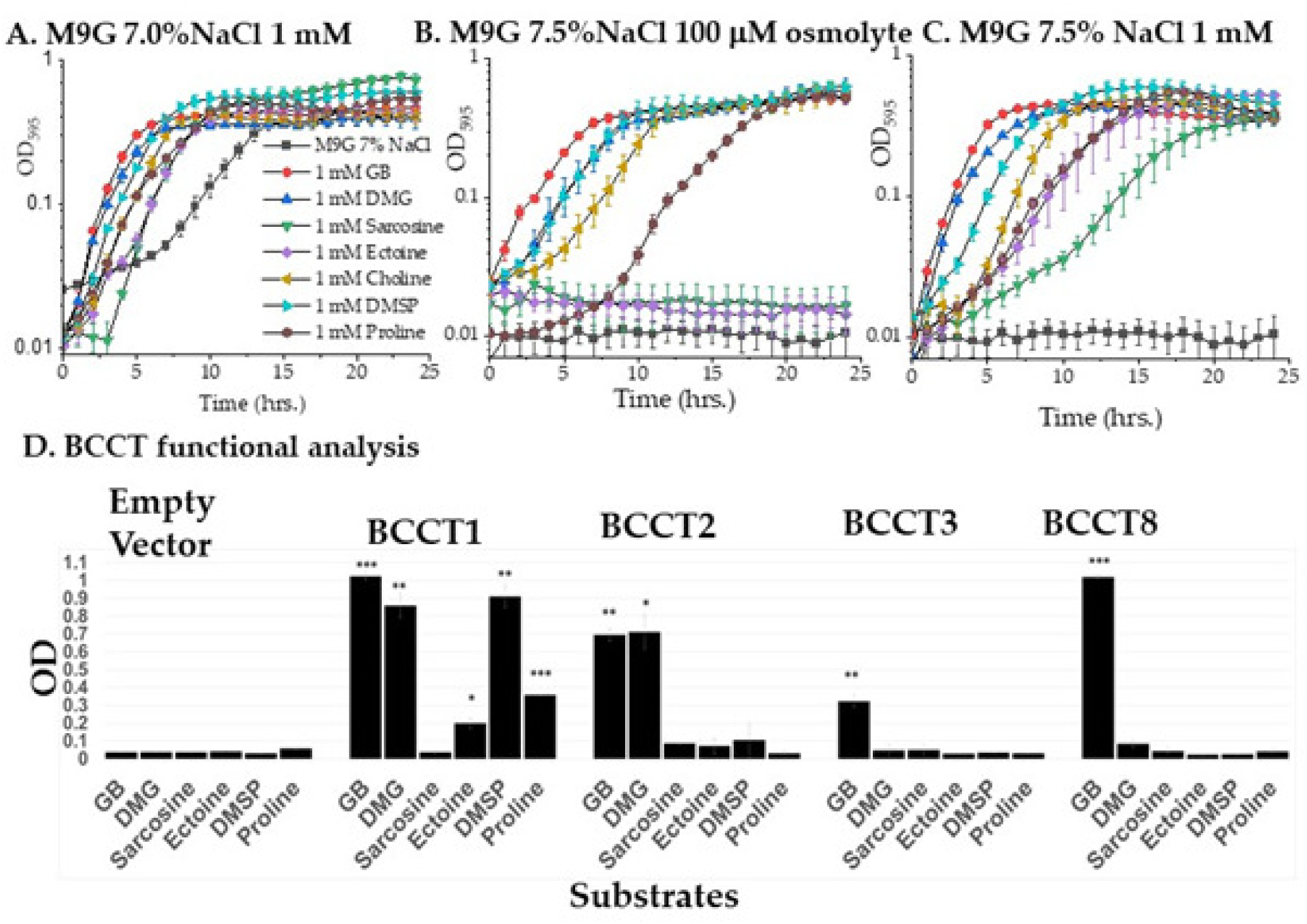
Import of compatible solutes for osmotic protection in *Vibrio natriegens* ATCC 14048. **A.** *V. natriegens* ATCC 14048 growth in M9G 7% NaCl with 1 mM GB (glycine betaine), DMG (dimethylglycine), sarcosine, ectoine, choline, DMSP (dimethylsulfoniopropionate) or L-proline. **B**. *V. natriegens* growth in M9G 7.5% NaCl in 100 µM GB, DMG, sarcosine, ectoine, choline, DMSP, or L-proline and **C.** 1 mM GB, DMG, sarcosine, ectoine, choline, DMSP, or L-proline. OD was measured for 24 h at 37°C**. D.** Functional complementation of *E. coli* MKH13 harboring *bccT1*, *bccT2*, *bccT3*, and *bccT8* grown in M9G–4% NaCl medium in the absence or presence of exogenously supplied compatible solutes GB, DMG, sarcosine, ectoine, DMSP or L-proline. The error bars represent means ± standard errors for two biological replicates. Statistical analysis was performed by comparing the *bccT* complemented *E. coli* strain to the corresponding empty vector supplemented with the same compatible solute. ****, P ≤ 0.001.

### BCCT1, BCCT2, BCCT3, and BCCT8 can transport osmolytes into the cell

Seven BCCTs were present in the genome of *V. natriegens,* and to determine the functionality of each, we cloned each *bccT* gene into *Escherichia coli* MKH13. The MKH13 strain of *E. coli* is a *putP, proP, proU,* and *betT-betIBA* deletion mutant, which prevents it from transporting osmolytes and converting choline into GB, which prevents growth in 4%NaCl (4). Functional complementation of *E. coli* MKH13 was performed by first cloning each *bccT* gene into the expression plasmid pBAD and then transforming each into *E. coli* MKH13. An empty vector plasmid was also cloned into *E. coli* MKH13 as a negative control and showed no growth in 4%NaCl in the presence of all osmolytes tested (**Fig. 4D**). *E. coli* pVn*bccT1*, pVn*bccT2*, pVn*bccT3*, and pVn*bccT8* were able to grow in the presence of GB, indicating these BCCTs transported GB into the cell for osmoprotection (**Fig. 4D**). *E. coli* pVn*bccT1* was able to uptake GB, DMG, DMSP, ectoine, and L-proline (**Fig. 4D**), and *E. coli* pVn*bccT2* transported GB, DMG, and DMSP (**Fig. 4D**). None of the transporters examined were able to uptake sarcosine, indicating a different transporter system is responsible (**Fig. 4D**).

Additionally, *E. coli* pVn*bccT4*, pVn*bccT6*, and pVn*bccT7* were unable to uptake any of the osmolytes tested (data not shown). Previous work by Gregory and colleagues characterized the GB binding pocket in BCCT1 from *V. parahaemolyticus* (31). BCCTs all contain 12 transmembrane (TM) domains, TM1 to TM12 and in *V. parahaemolyticus* BCCT1, the GB binding pocket residues were identified in TM4 and TM8. The specific amino acid identified as crucial for GB binding were Trp203, Trp208, and Tyr211 in TM4, and Trp380, Trp381, and Trp384 in TM8 (31). Our bioinformatics analysis of BCCT1, BCCT2, BCCT3, and BCCT8 from *V. natriegens* identified each of these residues in the corresponding TM4 and TM8 regions (**Fig. S2**). However, analysis of BCCT4, BCCT6, and BCCT7 showed a change in at least one of these essential residues (**Fig. S2**). For example, BCCT4, BCCT6, and BCCT7 had a change in the Tyr residue of the GB binding pocket in TM4, and BCCT4 and BCCT7 had a change in the 2nd and 3rd Trp residue of TM8 (**Fig. S2**). These changes to the GB binding pocket may be the cause of the inability of BCCT4, BCCT6, and BCCT7 to uptake GB. Since their expression and stability were not tested it is not possible to rule out their transport.

### *Vibrio natriegens* ATCC 14048 can utilize GB, DMG, and sarcosine as sole carbon sources

In *V. natriegens* ATCC 14048, the genomic context of *bcct8 (PN96_RS22655)* presence in chromosome 2 was examined. The *bcct8* gene formed a cluster, with the operon *dgcAB_fixAB (PN96_RS22675* to *PN96_RS22655),* four genes previously identified in *Pseudomonas aeruginosa* PAO1 (**Table S1) (Fig. 5)** (50). The *fixAB* genes were predicted to encode heterodimeric electron transfer flavoprotein alpha and beta subunits. The *dgcA* gene (*PN96_RS22675*) encodes a protein with a NADH:flavin oxidoreductase domain while *dgcB* (*PN96_RS22670*) encodes a protein with a Fe-S cluster containing an oxidoreductase domain and a domain of unknown function DUF3483 (**Table S1**). The *dgcAB* genes encode the alpha and beta subunits of a DMG dehydrogenase involved in the demethylation of DMG to sarcosine. Two additional genes preceded the *dgcAB_fixAB* cluster, encoding a hypothetical protein and a predicted dipeptidase (*pepD*). These two genes are also present at the same location in *P. aeruginosa* (**Fig. 5**) (50). Approximately 28-kb downstream of *pepD_hp_dgcAB_fixAB* was a second gene cluster (*PN96_RS22500* to *PN96_RS22455)* that contained genes encoding proteins required for catabolism of GB to DMG (*gbcA, gbcB*), sarcosine to pyruvate (*soxBDAG, glyA1, sdaB*), and the *purU* gene (**Fig. 5**). PurU is known to convert methyltetrahydrofolate (mTHF), a byproduct the *dgcAB* and *soxBDAG* enzymatic reactions, into tetrahydrofolate (THF) (**Table S1**) (48, 69). The region that separated the two GB catabolism clusters in *V. natriegens* contained gene homologs involved in the α-aminobutyrate (GABA) shunt pathway (*gabT, gabD*), amongst others (**Fig. 5**).

**Fig. 5.**
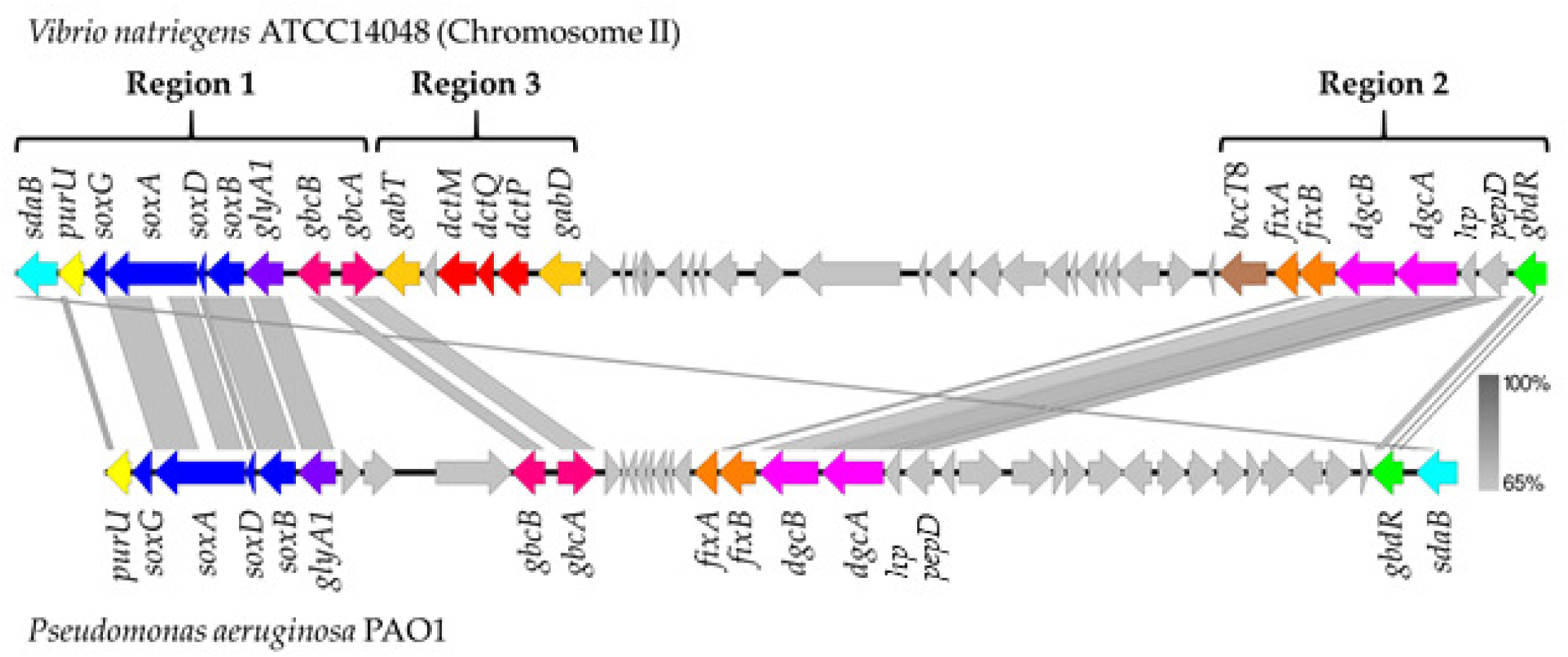
Genome comparison. Comparative analysis between *Vibrio natriegens* and *Pseudomonas aeruginosa* characterized GB catabolism genes. Arrows indicate open reading frames (ORFs) and the direction of transcription. Similarly colored arrows indicate genes with similar functions. Grey vertical bars indicate homology between genes.

To determine whether *V. natriegens* can grow on GB as a sole carbon source, growth was examined at 37°C in M9 supplemented with 3% NaCl or 1% NaCl with 20 mM GB as a sole carbon source. Under these conditions, no growth was observed (data not shown). Next, we examined growth at 30°C in M9 1% NaCl supplemented with 20 mM GB as sole carbon sources. Growth pattern analysis showed *V. natriegens* grew in GB with an approximately 1-h lag phase compared to M9G and a final OD of 0.22 (**Fig. 6A**). Next, we examined growth in DMG or sarcosine as sole carbon sources. In DMG, there was an approximately 1-h lag phase with a final OD of 0.38; whereas, on sarcosine there was a 4-h lag phase with a final OD of 0.53. *V. natriegens’* growth on sarcosine compared favorably with growth on glucose, which had a final OD of 0.61, indicating a similar biomass is produced on each substrate (**Fig. 6A**). Examination for growth on M9 1% NaCl supplemented with choline as a sole carbon source showed no growth (data not shown). To demonstrate the requirement for *gbcA* in the initial demethylation of GB into DMG in *V. natriegens*, we generated a *gbcA* non-polar in-frame deletion mutant by splice overlapping extension (SOE) PCR and allelic exchange. The *V. natriegens* Δ*gbcA* mutant was grown at 30°C in M9 1% NaCl supplemented with 20 mM glucose, GB, DMG, or sarcosine. Growth of the mutant in M9G was similar to wild-type indicating the mutant had no overall growth defect (**Fig. 6B**). The Δ*gbcA* mutant grew in media supplemented with DMG or sarcosine with an OD similar to growth on M9G but was unable to use GB as a sole carbon source. These data confirmed *gbcA* is required for GB catabolism but was not required for the breakdown of DMG and sarcosine, and the mutant does not exhibit any overall growth defect (**Fig. 6B**). This demonstrates that in *V. natriegens*, *gbcA* is only involved in the initial breakdown of GB into DMG confirming the first step of our putative catabolism pathway in *V. natriegens*.

**Fig. 6.**
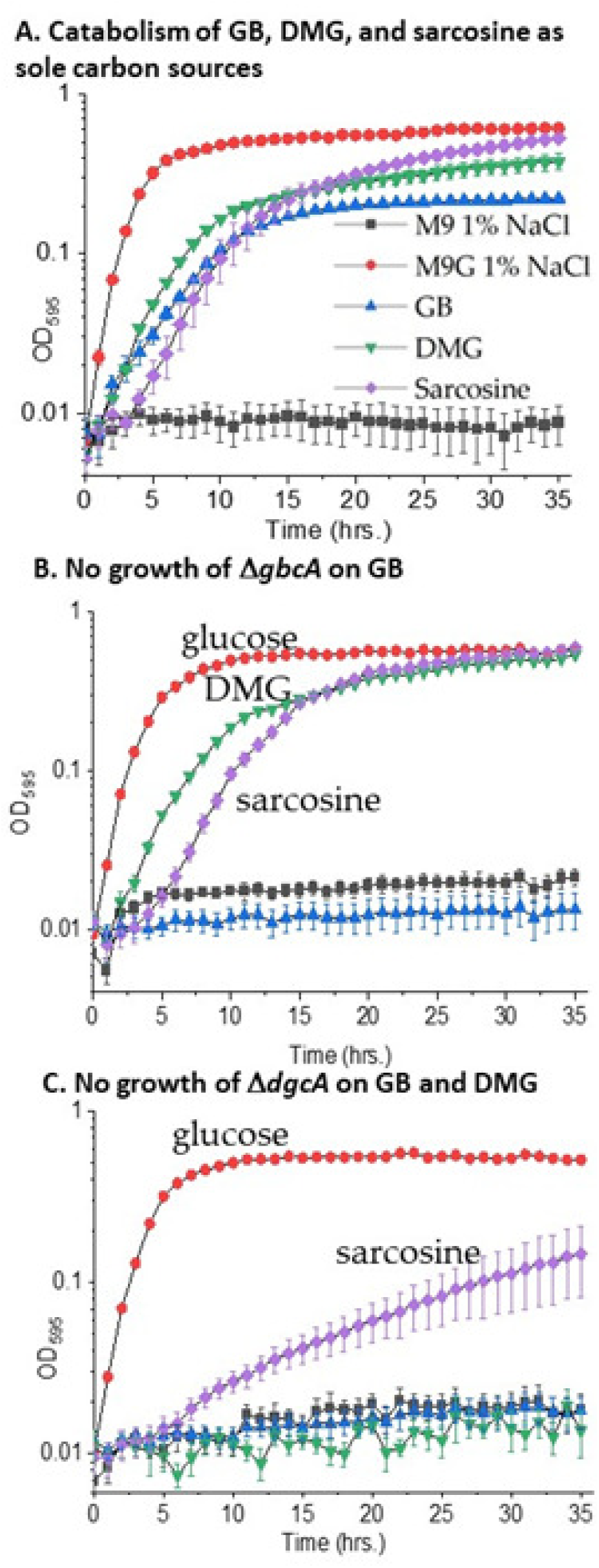
Utilization of GB, DMG, and sarcosine as sole carbon sources by *Vibrio natriegens*. **A.** Growth curves of *V. natriegens* ATCC 14048 in minimal media (M9) 1% NaCl at 30°C containing 20 mM glucose (M9G) or 20 mM glycine betaine (GB,), dimethylglycine (DMG,), sarcosine or no carbon source (M9) after 35 h. **B.** Growth curves of the *V. natriegens gbcA* deletion mutant in M9 1% NaCl at 30°C on glucose (M9G), GB, DMG, and sarcosine. **C.** Growth curves of the *V. natriegens dgcA* deletion mutant in M9 1% NaCl at 30°C in glucose (M9G), GB, DMG, and sarcosine.

The second step of GB catabolism, DMG degradation to sarcosine, was accomplished by the *dgcAB* genes in *P. aeruginosa*. In *V. natriegens*, a Δ*dgcA* mutant was constructed and examined for growth on M9G, M9 GB, M9 DMG, or M9 sarcosine. The Δ*dgcA* mutant grew on M9G similar to wild-type, showing no overall growth defect by the mutant (**Fig. 6C**). The mutant failed to grow in M9 GB or M9 DMG but grew on M9 sarcosine with a lower OD compared to M9G. This established that in *V. natriegens, dgcA* was involved in the breakdown of DMG to sarcosine, confirming the second step in the GB catabolism pathway.

### *Vibrio fluvialis* 2013V-1197 can utilize DMG and sarcosine as sole carbon sources

The catabolic genes were present as a single contiguous cluster among a subset of strains 39 strains) of *V. fluvialis,* an emerging seafood-borne pathogen that causes sporadic outbreaks of gastroenteritis worldwide (70). *Vibrio fluvialis* had a similar repertoire of osmotic stress response systems as other vibrios, containing the gene clusters for the biosynthesis of ectoine and GB as well as four BCCT transporters, BCCT1, BCCT2, BCCT3 and BCCT9 (no BCCT8 homolog was present) and two ProU transporters (**Fig. S3A**). *Vibrio fluvialis* 2013V-1197 showed growth in M9G 1% to 6% NaCl with growth prohibited in M9G 7% NaCl at 37°C (**Fig. S3B**). This species could uptake GB, DMG, DMSP, and ectoine to rescue growth in M9G 7% NaCl (**Fig. 7A**). Sarcosine did not rescue growth under the conditions examined (**Fig. 7A**).

**Fig. 7.**
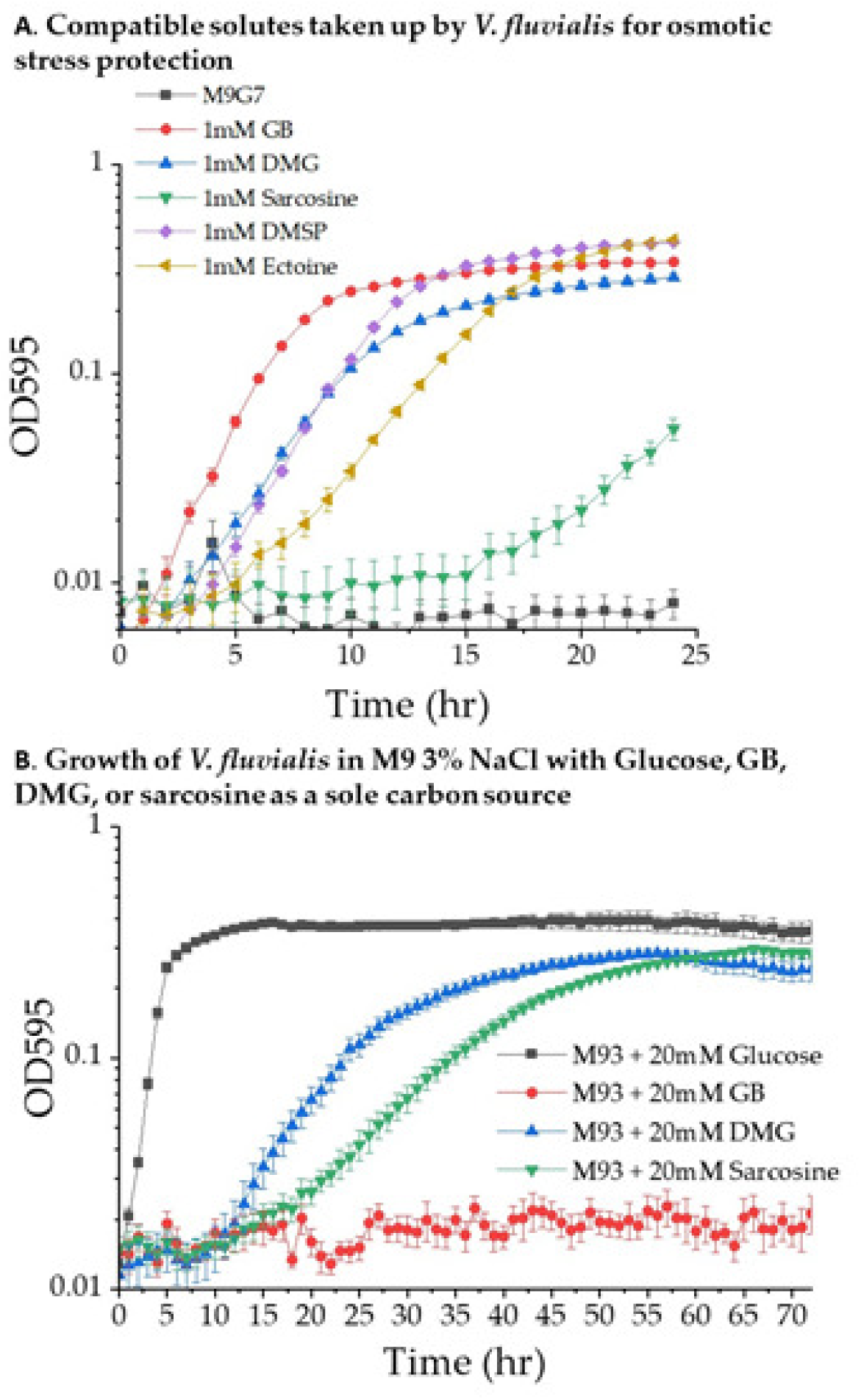
Compatible solutes as osmoprotectants and carbon sources in *V. fluvialis*. **A.** Import of compatible solutes for osmotic stress protection. *V. fluvialis* 2013V-1197 grown in M9G 7.0%NaCl supplemented with 1 mM GB, DMG, sarcosine, DMSP or ectoine. Growth was measured for 24 h at 37°C. **B.** Catabolism of compatible solutes. Growth curves of *V. fluvialis* 2013V-1197 in M9 3% NaCl at 37°C on glucose (M9G), GB, DMG, and sarcosine after 70h.

The ability of *V. fluvialis* 2013V-1197 to grow in M9 supplemented with GB, DMG, or sarcosine as sole carbon sources was examined in M9 1% NaCl at 30°C and 37°C. No growth was observed under these conditions (data not shown). Next, we examined growth in M9 3% NaCl at 30°C and 37°C supplemented with choline, GB, DMG, or sarcosine (**Fig. S4A and S4B**). No growth occurred in choline or GB as a sole carbon source after 72 h at 30°C or 37°C. At 30°C growth occurred in DMG or sarcosine after 48 h, whereas at 37°C growth occurred in DMG after 24 h (**Fig. S4A and S4B**).

Next, growth curve analyses were performed in M9 3% NaCl supplement with GB, DMG, or sarcosine. In M9 3% NaCl supplemented with DMG the growth curves showed a lag phase of 12 h with a final OD of 0.27 (**Fig. 7B**). In M9 3% NaCl supplemented with sarcosine with a lag phase of 14 h and a final OD of 0.27 after 60 h (**Fig. 7B**). Overall, these data show *V. fluvialis* 2013V-1197 cannot use GB as a sole carbon source but can consume either DMG or sarcosine as a sole carbon source.

### Phylogenetic analysis of DgcA and genome neighborhood analysis among marine bacteria

The GbcA and DgcA proteins among *V. natriegens* strains shared >99% identity and were present in all strains in chromosome 2 at the same location, but were present in two chromosomal regions (**Fig. 8**). Whereas among other *Vibrio* species, the catabolism genes was present as a contiguous cluster (**Fig. 8**). In *V. fluvialis* 2013V-1197, the r GB catabolism genes were present in chromosome 2, and in *V. gazogenes*, *V. mangrovi, V. ruber,* and *V. spartinae* strains, the region was present in chromosome 1 (**Fig. 8**). In *V. fluvialis* 2013V-1197, the region was flanked on either side by integrases inserted close to core genes, a D-alanine--D-alanine ligase on one side and H-NS on the other. An IS4 element was also noted near the GB catabolism region in *V. ruber,* and in *V. gazogenes* and *V. spartinae.* The region is flanked on one side by Rhs proteins and a few type VI secretion system genes and inserted adjacent to an asparagine-tRNA ligase in these species. This is also the insertion site for the region in *V. mangrovi*, but no integrases, IS elements or Rhs proteins were present. In all *V. penaeicida* strains, the GB biosynthesis genes (*betIBA*) and transporter genes (*proVWX/choVWX*) were directly upstream of the GB catabolic genes also in chromosome 1 (**Fig. 8**). In *V. ostrea* OG9-811, the *glyA1_soxBDAG_purU_sdaB* genes were present in chromosome 1, and *gbcA, gbcB, pepD*_*hp_dgcAB_fixAB* genes were present in chromosome 2. These data suggest independent acquisition of the region among species.

**Fig. 8.**
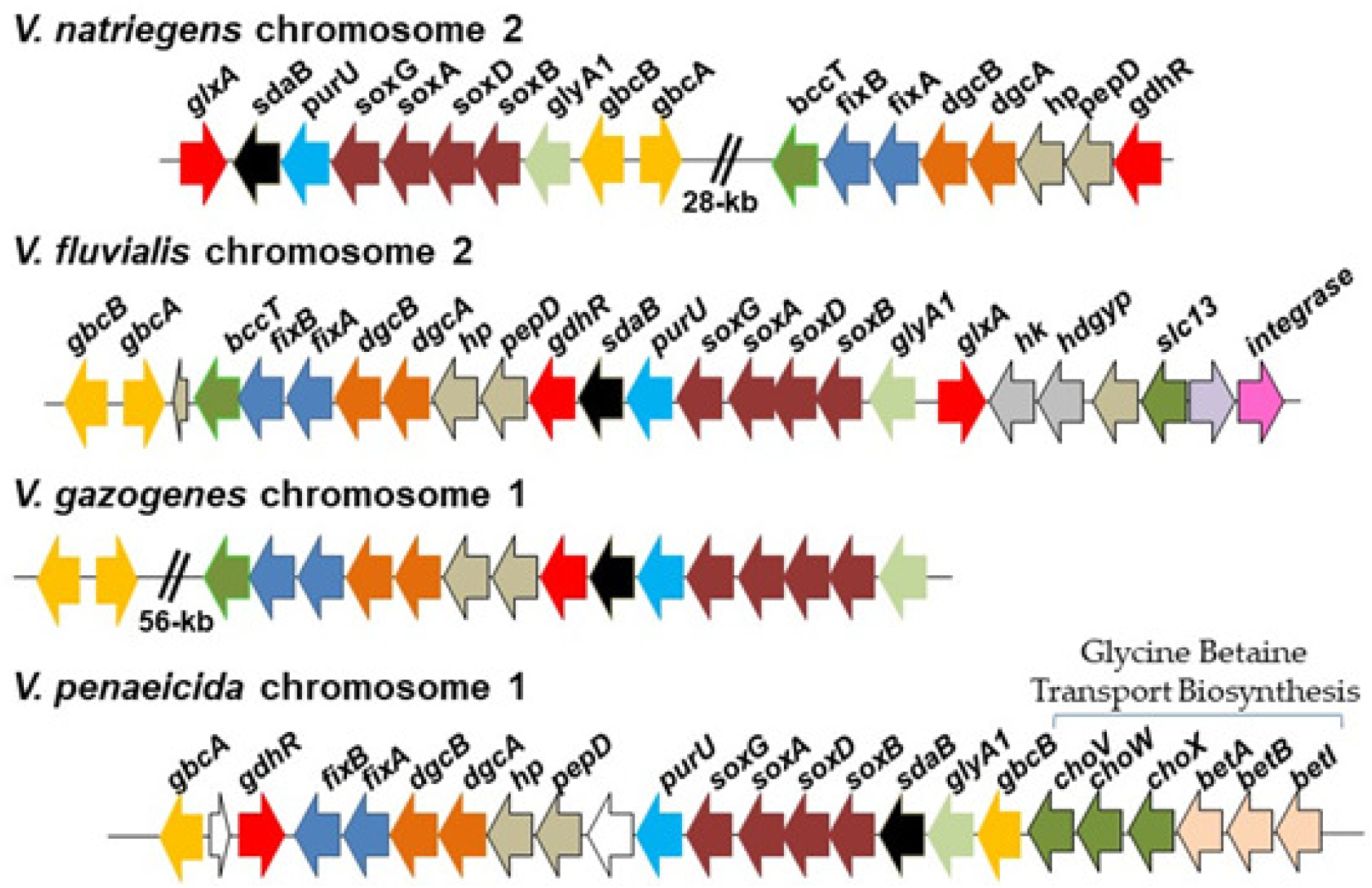
Schematic of the gene order of GB, DMG, and sarcosine transporter, catabolism and regulatory genes in *Vibrio* species. Arrows indicate ORFs and direction of transcription. Similarly colored arrows indicate genes sharing a similar function, with the exception of white arrows that represent genes of unknown function in most cases. In *V. penaeicida*, the GB biosynthesis genes (*betIBA*) and transporter (*choXWV*) genes are directly upstream of GB catabolism genes.

To further uncover the evolutionary history of GB, DMG, and sarcosine catabolism among *Vibrionaceae*, a phylogeny based on the DgcA protein was constructed (**Fig. 9**). The phylogeny placed DgcA from *Vibrionaceae* species within two highly divergent branches, with one branch comprised of *Vibrio* species sharing a most recent common ancestor several *Pantoea* and *Serratia* species (**Fig. 9**). In these two genera, the GB catabolism cluster contained a BCCT-type transporter similar to vibrios (**Fig. S5**). In *Serratia*, the cluster had a toxin-antitoxin system within the cluster and was directly downstream of a type I-C CRISPR-Cas system (**Fig. S5**). DgcA was present in most species of the genera *Marinobacterium*, *Marinomonas,* and *Marinobacter* amongst others. The second *Vibrionaceae* branch was comprised of *Grimontia, Enterovibrio* and *Photobacterium* clustered with DgcA from *V. penaeicida*. This was the most divergent branch on the tree (**Fig. 9**). These data show the catabolism region was acquired more than once within the *Vibrionaceae*. DgcA was present in all *Pseudomonas* species and a representative number are shown in the tree, which all clustered together indicating a common ancestor (**Fig. 9**).

**Fig. 9.**
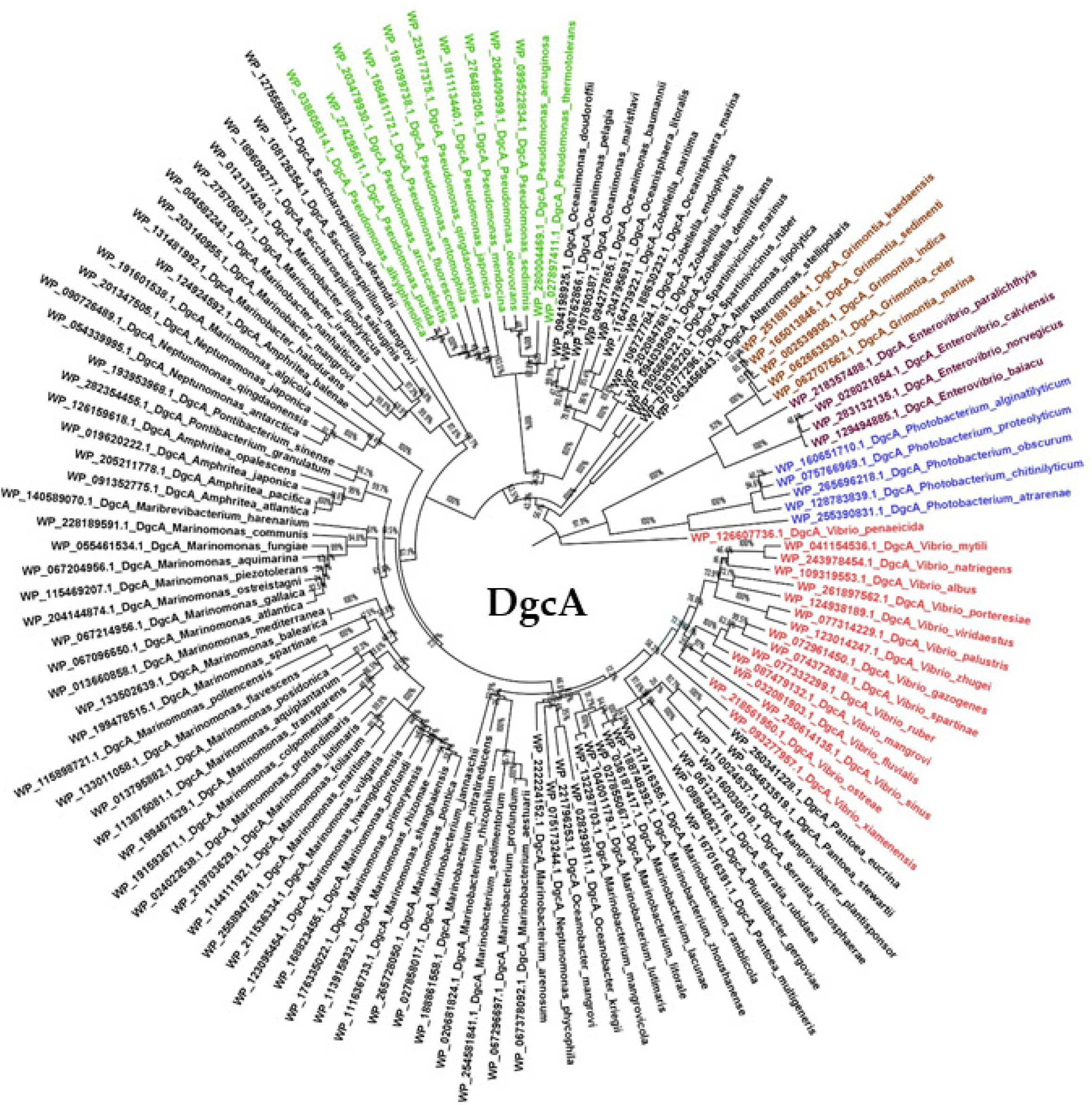
The evolutionary history of DgcA among *Vibrionaceae* using the Neighbor-Joining method with distances computed using the JTT matrix-based method and are in the units of the number of amino acid substitutions per site. The percentage of replicate trees in which the associated taxa clustered together in the bootstrap test (1000 replicates) are shown next to the branches. This analysis involved 126 amino acid sequences and ambiguous positions were removed using pairwise deletion option. Shown in red are *Vibrio* species, purple *Enterovibrio*, brown *Grimontia*, and blue *Photobacterium*, and green *Pseudomonas* species (only a subset of species from this genus is shown).

## DISCUSSION

*Vibrio natriegens* is a halophile and is non-pathogenic to humans, with a doubling time of less than 10 minutes under optimal growth conditions, which is proposed to be a function of its high substrate uptake rate (67, 68). Due to these attributes, there is a growing interest in using *V. natriegens* as an alternative cloning and expression tool in biotechnology (67, 71–78). Here, the osmotic stress adaptation systems were identified and characterized. We demonstrated that *V. natriegens* has high osmotic stress tolerance with an ability to grow in up to 7% NaCl without the addition of exogenous osmolytes. *Vibrio natriegens* was able to biosynthesize ectoine, glutamate, and glycine betaine (GB) as well as accumulate a variety of osmolytes utilizing BCCT transporters in response to osmotic stress. There were as many as four more BCCTs than are present in *V. parahaemolyticus* suggesting possibly wider substrate uptake abilities and/or different substrate affinities. At 37°C, *V. natriegens* was able to survive in 7% NaCl, but could not grow in the absence of NaCl. At 30°C, cells survived up to 6% NaCl and grew in the absence of NaCl. Addition of GB did not rescue growth at 37°C in the absence of NaCl. These data show that NaCl has a temperature-protective effect and that this protection is likely due to a decrease in water activity in the presence of NaCl. Choline, GB, DMG, and DMSP were shown to be highly effective osmolytes at both low and high osmolyte concentrations, rescuing growth in M9G 7.5% NaCl. GB, DMG, and DMSP were taken up efficiently by several BCCT transporters. In contrast, ectoine was only taken up by BCCT1 and sarcosine by none. In *V. natriegens*, BCCT1 took up the widest range of osmolytes, similar to what was shown for BCCT1 in *V. parahaemolyticus* (31). Previously, it was shown BCCT1, BCCT3, and BCCT4 in *V. parahaemolyticus* were highly induced under osmotic stress conditions (31). Whereas a recent study of *V. natriegens,* showed that the ABC-type ProU (aka *choXWV)* transporters were significantly induced in 300 mM NaCl (∼1.8% NaCl) compared to 0 mM, but none of the BCCT genes were induced (67).

Our data show that *V. natriegens* biosynthesized GB from choline and transported GB into the cell for osmotic stress protection at high NaCl concentrations, but unstressed cells did not. Unstressed *V. natriegens* cells grew on GB, DMG, and sarcosine as sole carbon and energy sources, but these cells could not use choline as a substrate to grow on. This suggests that choline is only directed to GB biosynthesis for osmoprotection, and osmolarity of the growth media dictated whether GB, DMG, and sarcosine are used as nutrients or for osmotic stress protection. Interestingly, temperature also played a role in the ability to catabolize these osmolytes in *V. natriegens*. The absence of growth on GB, DMG and sarcosine at 37°C even in 1% NaCl suggests that the catabolism and/or transporter genes may not be expressed at this temperature. In the rhizosphere associated bacterium *S. meliloti*, GB is a significant osmoprotectant with uptake strongly induced under osmotic stress (43–46). GB and choline catabolism was inhibited in *S. meliloti* stressed cells, but growth on GB and choline was highly active in unstressed cells, demonstrating the importance of GB for stress protection over catabolism (43–46). This is in contrast to *P. aeruginosa*, a species that catabolizes both choline and GB in relatively high NaCl concentrations (0.75 M NaCl, which equals approx. 4.25% NaCl). In *P. aeruginosa* and other Pseudomonads, it was shown that choline and GB intracellular pools are maintained under a range of growth conditions (79). In *Chomohalobacter salexigens*, only GB supported growth as a sole carbon source and only when cells were grown under high NaCl concentrations (58, 59). Our work showed that in *V. fluvialis,* DMG and sarcosine supported growth in M9 3% NaCl at 30°C and 37°C, but choline and GB did not. In addition, DMG and sarcosine had longer lag phases and lower ODs compared to *V. natriegens* showing less efficient utilization of these substrates. We speculate that there must be significant differences in the regulation of GB, DMG, and sarcosine catabolism and/or transporter genes between these species, which will require further examination.

Analysis of the evolutionary history of the catabolism genes was revealing as it demonstrated the presence of the gene cluster with very different evolutionary histories. Within *Vibrionaceae*, the region was present in a limited number of species and was acquired more than once, reflected by its presence on different chromosomes, different gene arrangements, and different evolutionary histories. Among *Marinobacterium* and *Marinomonas* species DgcA was prevalent with the entire cluster present in these genera suggesting a functional GB pathway. We speculate that acquisition of the catabolism genes provided an evolutionary advantage allowing a species a competitive advantage in specific environments such as on and within plant tissues, as animal pathogens or associated with marine flora and fauna.

## MATERIALS AND METHODS

### Bacterial strains, media, and growth conditions

All bacterial strains and plasmids used in this study are listed in **Table S2.** *V. natriegens* ATCC 14048 and *V. fluvialis* 2013V-1197 were used as wild-type (WT). Strains was grown in either lysogeny broth (LB) (NaCl, 5 g/L, tryptone, 10 g/L, yeast extract, 5 g/L) (Thermo Fisher Scientific, Fair Lawn, NJ) final 3% NaCl (wt/vol) (LB 3% NaCl) or M9 minimal media (47.8 mM Na_2_HPO_4_, 22 mM KH_2_PO_4_, 18.7 mM NH_2_Cl, and 8.6 mM NaCl; Sigma-Aldrich) supplemented with 2 mM MgSO_4_, 0.1 mM CaCl_2_, and 20 mM D-glucose (Sigma-Aldrich) (M9G) as the sole carbon source and NaCl as indicated. Choline, GB, DMG, and sarcosine were used at a final concentration of 20 mM when used as a sole carbon source. *Escherichia coli* strains were grown in either LB supplemented with 1% NaCl (LB 1% NaCl) or M9G supplemented with 4% NaCl, as indicated. Media was supplemented with 0.3 mM diaminopimelic acid (DAP) (Sigma-Aldrich) and/or chloramphenicol (Cm) at 25 µg/mL when needed. The osmolytes GB, DMG, choline, ectoine, and DMSP were used at a final concentration of 100 µM or 1 mM. All strains were grown at either 37°C or 30°C, as indicated.

### Growth pattern analysis at varying NaCl concentrations and temperatures in M9 media in the absence and presence of compatible solutes

For growth analysis in varying salt concentrations, *V. natriegens* was grown overnight in M9G 2% NaCl at 37°C. A 2% inoculation of the overnight culture was grown to OD ∼0.55 at 37°C in fresh M9G 2% NaCl. Cultures were pelleted and washed twice in M9G to remove excess salt. A 1:40 dilution into 200 µL of M9G and M9G 1% to 7.5% NaCl was made in a 96-well plate. A Tecan Sunrise microplate reader was used to incubate the plates at 37°C or 30°C with intermittent shaking and to measure the optical density at 595 nm (OD_595_) hourly for 24 h. To examine the osmolytes used by V*. natriegens*, a 1:40 dilution was inoculated into either M9G 7% NaCl or M9G 7.5% NaCl supplemented with either 100µM or 1 mM of choline, GB, DMG, sarcosine, ectoine, DMSP, or L-proline and the OD_595_ was measured hourly for 24 h at 37°C.

### 1H-Nuclear magnetic resonance (^1^H-NMR) analysis of ectoine and GB biosynthesis

For analysis of ectoine and GB biosynthesis, *V. natriegens* ATCC 14048 was grown in M9G 1% NaCl, M9G 5% NaCl, or M9G 5 % NaCl supplemented with 1 mM choline at 37°C to stationary phase. For analysis of ectoine production, no choline was added to the medium. Cultures were pelleted by centrifugation for 10 min at 6,000 rpm. The supernatant was discarded, and the pellet was washed once with an isotonic solution. Cultures supplemented with choline were washed with an isotonic solution without choline. Cultures were pelleted by centrifugation for 10 min at 6,000 rpm, and the supernatant was discarded. Cells were lysed by three freeze-thaw cycles (1 h at −80°C then 30 min on ice). Pellets were suspended in 750 µL 190 proof ethanol and vortexed. The suspended solution was centrifuged for 10 min at 12,000 rpm to remove debris. The ethanol extract was evaporated by vacuum for 3.5 h. The extract was suspended in 700 µL deuterium oxide (D_2_O) and vortexed. Debris was removed by centrifugation for 10 min at 12,000 rpm. Extract was filter-sterilized to remove any debris not removed by centrifugation. The extract was transferred to a 5-mm NMR tube. Samples were analyzed in an AVIII 600 MHz NMR spectrometer’s qProton method with 16 scans per spectra. Data was analyzed using MestReNova (Mnova) NMR software.

### Functional complementation of the *E. coli* MKH13 strain with *bccT* genes

Genomic DNA from *V. natriegens* ATCC 14048 RefSeq GCF_001456255.1 (chromosome I: NZ_CP009977.1 and chromosome II: NZ_CP009978.1) was used as template to amplify *bccT1, bccT2, bccT3, bccT4, bccT6, bccT7,* and *bccT8* using the primers listed in **Table S3** and designed from RefSeq <u>NZ_CP009977.1</u> and <u>NZ_CP009978.1</u>. Primers were purchased from Integrated DNA Technologies (Coralville, IA) and each gene was PCR amplified and cloned into a pBAD 33 expression vector (80) via Gibson assembly, using NEBuilder HiFi DNA assembly master mix. Plasmids pBAVn*bccT*1, pBAVn*bccT*2, pBAVn*bccT*3, pBAVn*bccT*4, pBAVn*bccT*6, pBAVn*bccT*7, or pBAVn*bccT*8 were transformed and propagated in the *E. coli* DH5α strain (**Table S2**). The expression plasmids were purified and sequenced. The plasmids were then transformed into competent *E. coli* MKH13 cells (4)(**Table S2**). This strain contains deletions including the *betT*-*betIBA* genes for choline uptake and GB biosynthesis and the *putP*, *proP*, and *proU* genes required for osmolyte uptake. These deletions prevent MKH13 growth in 4% NaCl. The *E. coli* pVn*bccT*1, pVn*bccT*2, pVn*bccT*3, pVn*bccT*4, pVn*bccT*6, pVn*bccT*7, or pVn*bccT*8 strains were grown overnight in M9G 1% NaCl supplemented with 0.01% Cm, to maintain the plasmid. A 1:100 dilution was made of the overnight cultures into M9G 4% NaCl supplemented with 0.01% Cm and 0.01% L-arabinose, the inducing agent, and 1 mM of GB, DMG, sarcosine, ectoine, DMSP, or L-proline. Cultures were incubated at 37°C for 24 h and OD_595_ measured. Statistics were calculated using the t-Test: Two-Sample Assuming Equal Variances and compared to growth with pBAD33 empty vector.

### Growth pattern analysis of *V. natriegens* and *V. fluvialis* on choline, GB, DMG, and sarcosine as sole carbon sources

*V. natriegens* ATCC 14048 and *V. fluvialis* 2013V-1197 growth on choline, GB, DMG, and sarcosine as sole carbon sources was examined at 37°C and 30°C in M9 supplemented with 3% NaCl or 1% NaCl with 20 mM of each carbon source. *V. natriegens* was grown overnight in M9 3% NaCl and 20 mM glucose at 37°C. This cultures was used to generate a 2% inoculation in M9 media and grown to OD ∼ 0.55 at 37°C and then washed twice with PBS to remove glucose. Washed cells were inoculated 1:40 into 200 µL M9 media supplemented with 3% NaCl and 1% NaCl, and 20 mM of each carbon source. Plates were incubated at 37°C and 30°C with intermittent shaking for 1 min every hour. The optical densities were measured at 595 nm every hour for 35 h for *V. natriegens* and 60 h for *V. fluvialis* using a Tecan Sunrise microplate reader and Magellan plate reader software (Tecan Systems Inc., San Jose, CA).

### Catabolism gene deletion construction

In-frame non-polar gene deletions were constructed by SOE (splicing by overlap extension) PCR and homologous recombination (81). The *V. natriegens* ATCC 14048 genome sequence RefSeq GCF_001456255.1 (Chromosome II: NZ_CP009978.1) was used to generate fragments AB and CD, for each gene using the primers listed in **Table S3**. For *gbcA*, Gibson assembly was used to ligate the two fragments with a SacI digested suicide vector, pDS132 (82), to produce a vector with the truncated gene, pDS132VnΔ*gbcA*. pDS132VnΔ*gbcA* was transformed into the *E. coli* strain DH5α, with Cm, then into the DAP auxotroph *E. coli* strain β2155 λ*pir* (83), with Cm and DAP. This strain was used to perform conjugation with *V. natriegens*. The vector requires the *pir* gene in order to replicate. In order for pDS132ΔVn*gbcA* to replicate in *V. natriegens*, which lacks the *pir* gene, it must incorporate *pir* into the genome by homologous recombination. Colonies were grown with Cm and screened for the presence of the incorporated vector. To induce the second homologous recombination in which the truncated *gbcA* gene replaces wild-type *gbcA* in the *V. natriegens* genome, colonies were grown without Cm and in the presence of sucrose. The pDS132 vector contains the *sacB* gene which codes for levansucrase, which will form a toxic polymer with sucrose causing colonies that contain the plasmid to have an altered morphology with a soupy appearance. Colonies with a normal morphology were screened for the presence of the truncated *gbcA* gene. An identical protocol was followed for construction of a *dgcA* deletion mutant. In-frame deletions were confirmed by DNA sequencing.

### Phylogenetic analysis and gene neighbor analysis

BLAST analyses were used to identify homologous protein sequences in the NCBI genome database within and outside the *Vibrionaceae* family. Proteins with >95% query coverage and >60% amino acid similarity were downloaded and aligned using CLUSTALW (84). Phylogenetic trees were constructed from 126 DgcA proteins using the program MEGA11 (85). Protein sequences were downloaded from NCBI and aligned using CLUSTALW. Trees were generated using evolutionary distances computed by the JTT matrix-based method with rate variation modelled with a gamma distribution. The pairwise deletion option was applied to all ambiguous positions for each sequence pair resulting in a final data set comprising 698 positions.

## Data Availability

All datasets are available upon request.

## ACKNOWLEDGMENTS

K.E.B.L. was funded in part by the Chemistry-Biology Interface Predoctoral Training Program (grant 5T32GM008550). We thank Dr. Cheryl Tarr, CDC, Atlanta, GA, USA for the gift of the *V. fluvialis* strain 2013V-1197.

## REFERENCES

1. Galinski E, Oren A. 1991. Isolation and structure determination of a novel compatible solute from the moderately halophilic purple sulfur bacterium *Ectothiorhodospira marismortui*. Eur J Biochem 198:593–8.

2. Galinski EA. 1995. Osmoadaptation in bacteria. Adv Microb Physiol 37:272–328.

3. Boch J, Kempf B, Bremer E. 1994. Osmoregulation in *Bacillus subtilis*: synthesis of the osmoprotectant glycine betaine from exogenously provided choline. J Bacteriol 176:5364–71.

4. Haardt M, Kempf B, Faatz E, Bremer E. 1995. The osmoprotectant proline betaine is a major substrate for the binding-protein-dependent transport system ProU of *Escherichia coli* K-12. Mol Gen Genet 246:783–6.

5. da Costa MS, Santos H, Galinski EA. 1998. An overview of the role and diversity of compatible solutes in Bacteria and Archaea. Adv Biochem Eng Biotechnol 61:117–53.

6. Kunte H, Crane R, Culham D, Richmond D, Wood J. 1999. Protein ProQ influences osmotic activation of compatible solute transporter ProP in *Escherichia coli* K-12. J Bacteriol 181:1537–43.

7. Sleator RD, Hill C. 2002. Bacterial osmoadaptation: the role of osmolytes in bacterial stress and virulence. FEMS Microbiol Rev 26:49–71.

8. Wood JM. 2011. Bacterial osmoregulation: a paradigm for the study of cellular homeostasis. Annu Rev Microbiol 65:215–38.

9. Wood JM. 2015. Bacterial responses to osmotic challenges. J Gen Physiol 145:381–8.

10. Bremer E, Kramer R. 2019. Responses of microorganisms to osmotic stress. Annu Rev Microbiol 73:313–334.

11. Ongagna-Yhombi SY, Boyd EF. 2013. Biosynthesis of the osmoprotectant ectoine, but not glycine betaine, is critical for survival of osmotically stressed *Vibrio parahaemolyticus* cells. Appl Environ Microbiol 79:5038–49.

12. Czech L, Hermann L, Stoveken N, Richter AA, Hoppner A, Smits SHJ, Heider J, Bremer E. 2018. Role of the extremolytes ectoine and hydroxyectoine as stress protectants and nutrients: genetics, phylogenomics, biochemistry, and structural Analysis. Genes (Basel) 9.

13. Hermann L, Mais, C. N., Czech, Smits, S. H., Bange, G., Bremer,E. 2020. The ups and downs of ectoine: structural enzymology of a major microbial stress protectant and versatile nutrient. Biological Chemistry 401:1443–1468.

14. Kappes RM, Kempf B, Kneip S, Boch J, Gade J, Meier-Wagner J, Bremer E. 1999. Two evolutionarily closely related ABC transporters mediate the uptake of choline for synthesis of the osmoprotectant glycine betaine in *Bacillus subtilis*. Mol Microbiol 32:203–16.

15. Saier MH, Jr. 2000. Families of transmembrane transporters selective for amino acids and their derivatives. Microbiology 146 (Pt 8):1775–1795.

16. Holtmann G, Bremer E. 2004. Thermoprotection of *Bacillus subtilis* by exogenously provided glycine betaine and structurally related compatible solutes: involvement of Opu transporters. J Bacteriol 186:1683–93.

17. Malek AA, Chen C, Wargo MJ, Beattie GA, Hogan DA. 2011. Roles of three transporters, CbcXWV, BetT1, and BetT3, in Pseudomonas aeruginosa choline uptake for catabolism. J Bacteriol 193:3033–41.

18. Wood JM. 2007. Bacterial osmosensing transporters. Methods Enzymol 428:77–107.

19. Perez C, Koshy C, Ressl S, Nicklisch S, Kramer R, Ziegler C. 2011. Substrate specificity and ion coupling in the Na+/betaine symporter BetP. EMBO J 30:1221–9.

20. Ziegler C, Bremer, E., Kramer, R. 2010. The BCCT family of carriers: from physiology to crystal structure. Mol Microbiol 78:13–34.

21. Ongagna-Yhombi SY, McDonald ND, Boyd EF. 2015. Deciphering the role of multiple betaine-carnitine-choline transporters in the halophile *Vibrio parahaemolyticus*. Appl Environ Microbiol 81:351–63.

22. Teichmann L, Chen C, Hoffmann T, Smits SHJ, Schmitt L, Bremer E. 2017. From substrate specificity to promiscuity: hybrid ABC transporters for osmoprotectants. Mol Microbiol 104:761–780.

23. Hoffmann T, Bremer E. 2017. Guardians in a stressful world: the Opu family of compatible solute transporters from *Bacillus subtilis*. Biol Chem 398:193–214.

24. Breisch J, Averhoff B. 2020. Identification of osmo-dependent and osmo-independent betaine-choline-carnitine transporters in *Acinetobacter baumannii*: role in osmostress protection and metabolic adaptation. Environ Microbiol 22:2724–2735.

25. Naughton LM, Blumerman SL, Carlberg M, Boyd EF. 2009. Osmoadaptation among *Vibrio* species and unique genomic features and physiological responses of *Vibrio parahaemolyticus*. Appl Environ Microbiol 75:2802–10.

26. Wood JM, Bremer E, Csonka LN, Kraemer R, Poolman B, van der Heide T, Smith LT. 2001. Osmosensing and osmoregulatory compatible solute accumulation by bacteria. Comp Biochem Physiol A Mol Integr Physiol 130:437–60.

27. Hoffmann T, Wensing A, Brosius M, Steil L, Volker U, Bremer E. 2013. Osmotic control of opuA expression in *Bacillus subtilis* and its modulation in response to intracellular glycine betaine and proline pools. J Bacteriol 195:510–22.

28. Boas Lichty KE, Loughran RM, Ushijima B, Richards GP, Boyd EF. 2024. Osmotic stress response of the coral and oyster pathogen Vibrio coralliilyticus: acquisition of catabolism gene clusters for the compatible solute and signaling molecule myo-inositol. bioRxiv doi:10.1101/2024.01.16.575920:2024.01.16.575920.

29. Gregory GJ, Boas KE, Boyd EF. 2021. The organosulfur compound dimethylsulfoniopropionate (DMSP) is utilized as an osmoprotectant by *Vibrio* species. Appl Environ Microbiol 87:02235–20.

30. Gregory GJ, Boyd EF. 2021. Stressed out: Bacterial response to high salinity using compatible solute biosynthesis and uptake systems, lessons from *Vibrionaceae*. Computational and Structural Biotechnology Journal 19:1014–1027.

31. Gregory GJ, Dutta A, Parashar VJ, Boyd EF. 2020. Investigations of dimethylglycine (DMG), glycine betaine and ectoine uptake by a BCCT family transporter with broad substrate specificty in Vibrio species. J Bact 202:e00314–20.

32. Kapfhammer D, Karatan E, Pflughoeft KJ, Watnick PI. 2005. Role for glycine betaine transport in Vibrio cholerae osmoadaptation and biofilm formation within microbial communities. Appl Environ Microbiol 71:3840–7.

33. Reen FJ, Almagro-Moreno S, Ussery D, Boyd EF. 2006. The genomic code: inferring Vibrionaceae niche specialization. Nat Rev Microbiol 4:697–704.

34. McParland EL, Alexander H, Johnson WM. 2021. The osmolyte ties that bind: genomic insights into synthesis and breakdown of organic osmolytes in marine microbes. Frontiers in Marine Science 8.

35. Kiene RP. 1998. Uptake of choline and its conversion to glycine betaine by bacteria in estuarine waters. Appl Environ Microbiol 64:1045–51.

36. Kiene RP, Hoffmann Williams LP. 1998. Glycine betaine uptake, retention, and degradation by microorganisms in seawater. Limnology and Oceanography 43:1592–1603.

37. Kiene RP, L. P. Hoffmann, Walker. JE. 1998. Sea water microorganisms have a high-afnity glycine betaine uptake system which also recognizes dimethylsulfoniopropionate. . Aquat Microbiol Ecol 15:39–51.

38. Keller MD, Kiene RP, Matrai PA, Bellows WK. 1999. Production of glycine betaine and dimethylsulfoniopropionate in marine phytoplankton. I. Batch cultures. Marine Biology 135:237–248.

39. Ngugi DK, Ziegler M, Duarte CM, Voolstra CR. 2020. Genomic blueprint of glycine betaine metabolism in coral metaorganisms and their contribution to reef nitrogen budgets. iScience 23:101120.

40. Keller MD, Matrai PA, Kiene RP, Bellows WK. 2004. Responses of coastal phytoplankton populations to nitrogen additions: dynamics of cell-associated dimethylsulfoniopropionate (DMSP), glycine betaine (GBT), and homarine. Canadian Journal of Fisheries and Aquatic Sciences 61:685–699.

41. Airs RL, Archer SD. 2010. Analysis of glycine betaine and choline in seawater particulates by liquid chromatography/electrospray ionization/mass spectrometry. Limnology and Oceanography: Methods 8:499–506.

42. Kortstee GJJ. 1970. The aerobic decomposition of choline by microorganisms. Archiv für Mikrobiologie 71:235–244.

43. Bernard T, Pocard J-A, Perround B, Le Rudulier D. 1986. Variations in the response of salt-stressed Rhizobium strains to betaines. Archives of Microbiology 143:359–364.

44. Smith LT, Pocard JA, Bernard T, Le Rudulier D. 1988. Osmotic control of glycine betaine biosynthesis and degradation in *Rhizobium meliloti*. J Bacteriol 170:3142–9.

45. Goldmann A, Boivin C, Fleury V, Message B, Lecoeur L, Maille M, Tepfer D. 1991. Betaine use by rhizosphere bacteria: genes essential for trigonelline, stachydrine, and carnitine catabolism in Rhizobium meliloti are located on pSym in the symbiotic region. Mol Plant Microbe Interact 4:571–8.

46. Talibart R, Jebbar M, Gouffi K, Pichereau V, Gouesbet G, Blanco C, Bernard T, Pocard J. 1997. Transient accumulation of glycine betaine and dynamics of endogenous osmolytes in salt-stressed cultures of *Sinorhizobium meliloti*. Appl Environ Microbiol 63:4657–63.

47. Barra L, Fontenelle C, Ermel G, Trautwetter A, Walker GC, Blanco C. 2006. Interrelations between glycine betaine catabolism and methionine biosynthesis in *Sinorhizobium meliloti* strain 102F34. J Bacteriol 188:7195–204.

48. Meskys R, Harris RJ, Casaite V, Basran J, Scrutton NS. 2001. Organization of the genes involved in dimethylglycine and sarcosine degradation in *Arthrobacter* spp. European Journal of Biochemistry 268:3390–3398.

49. Diab F, Bernard T, Bazire A, Haras D, Blanco C, Jebbar M. 2006. Succinate-mediated catabolite repression control on the production of glycine betaine catabolic enzymes in *Pseudomonas aeruginosa* PAO1 under low and elevated salinities. Microbiology (Reading) 152:1395–1406.

50. Wargo MJ, Szwergold BS, Hogan DA. 2008. Identification of two gene clusters and a transcriptional regulator required for *Pseudomonas aeruginosa* glycine betaine catabolism. J Bacteriol 190:2690–9.

51. Wargo MJ. 2013. Choline catabolism to glycine betaine contributes to *Pseudomonas aeruginosa* survival during murine lung infection. PLoS One 8:e56850.

52. Wargo MJ. 2013. Homeostasis and catabolism of choline and glycine betaine: lessons from *Pseudomonas aeruginosa*. Appl Environ Microbiol 79:2112–20.

53. Cánovas D, Vargas C, Csonka LN, Ventosa A, Nieto JJ. 1996. Osmoprotectants in *Halomonas elongata*: high-affinity betaine transport system and choline-betaine pathway. J Bacteriol 178:7221–6.

54. Vargas C, Jebbar M, Carrasco R, Blanco C, Calderón MI, Iglesias-Guerra F, Nieto JJ. 2006. Ectoines as compatible solutes and carbon and energy sources for the halophilic bacterium *Chromohalobacter salexigens*. J Appl Microbiol 100:98–107.

55. Wargo MJ, Ho TC, Gross MJ, Whittaker LA, Hogan DA. 2009. GbdR regulates *Pseudomonas aeruginosa plcH* and *pchP* transcription in response to choline catabolites. Infect Immun 77:1103–11.

56. Hampel KJ, LaBauve AE, Meadows JA, Fitzsimmons LF, Nock AM, Wargo MJ. 2014. Characterization of the GbdR regulon in *Pseudomonas aeruginosa*. J Bacteriol 196:7–15.

57. Willsey GG, Wargo MJ. 2016. Sarcosine catabolism in *Pseudomonas aeruginosa* is transcriptionally regulated by SouR. J Bacteriol 198:301–10.

58. Shao YH, Guo LZ, Zhang YQ, Yu H, Zhao BS, Pang HQ, Lu WD. 2018. Glycine Betaine monooxygenase, an unusual rieske-type oxygenase system, catalyzes the oxidative N-demethylation of glycine betaine in *Chromohalobacter salexigens* DSM 3043. Appl Environ Microbiol 84.

59. Yang T, Shao YH, Guo LZ, Meng XL, Yu H, Lu WD. 2020. Role of N,N-Dimethylglycine and its catabolism to sarcosine in Chromohalobacter salexigens DSM 3043. Appl Environ Microbiol 86.

60. Pichereau V, Pocard JA, Hamelin J, Blanco C, Bernard T. 1998. Differential effects of dimethylsulfoniopropionate, dimethylsulfonioacetate, and other S-methylated compounds on the growth of *Sinorhizobium meliloti* at low and high osmolarities. Appl Environ Microbiol 64:1420–9.

61. Bremer E. 2014. Liberate and grab it, ingest and digest it: the GbdR regulon of the pathogen *Pseudomonas aeruginosa*. J Bacteriol 196:3–6.

62. Xu J, Chen Q, Lønborg C, Li Y, Cai R, He C, Shi Q, Hu Y, Wang Y, Jiao N, Zheng Q. 2022. You exude what you eat: How Carbon-, Nitrogen-, and Sulfur-rich organic substrates shape microbial community composition and the dissolved organic matter pool. Applied and Environmental Microbiology 88:e01558–22.

63. Boysen AK, Carlson LT, Durham BP, Groussman RD, Aylward FO, Ribalet F, Heal KR, White AE, DeLong EF, Armbrust EV, Ingalls AE. 2021. Particulate Metabolites and Transcripts Reflect Diel Oscillations of Microbial Activity in the Surface Ocean. mSystems 6.

64. Boysen AK, Durham BP, Kumler W, Key RS, Heal KR, Carlson LT, Groussman RD, Armbrust EV, Ingalls AE. 2022. Glycine betaine uptake and metabolism in marine microbial communities. Environmental Microbiology 24:2380–2403.

65. Mausz MA, Airs RL, Dixon JL, Widdicombe CE, Tarran GA, Polimene L, Dashfield S, Beale R, Scanlan DJ, Chen Y. 2022. Microbial uptake dynamics of choline and glycine betaine in coastal seawater. Limnology and Oceanography 67:1052–1064.

66. Payne WJ, Eagon RG, Williams AK. 1961. Some observations on the physiology of *Pseudomonas natriegens* nov. spec. Antonie van Leeuwenhoek 27:121–128.

67. Coppens L, Tschirhart T, Leary DH, Colston SM, Compton JR, Hervey WJt, Dana KL, Vora GJ, Bordel S, Ledesma-Amaro R. 2023. *Vibrio natriegens* genome-scale modeling reveals insights into halophilic adaptations and resource allocation. Mol Syst Biol 19:e10523.

68. Hoffart E, Grenz S, Lange J, Nitschel R, Müller F, Schwentner A, Feith A, Lenfers-Lücker M, Takors R, Blombach B. 2017. High substrate uptake rates empower *Vibrio natriegens* as production host for industrial biotechnology. Appl Environ Microbiol 83.

69. Lidbury I, Kimberley G, Scanlan DJ, Murrell JC, Chen Y. 2015. Comparative genomics and mutagenesis analyses of choline metabolism in the marine Roseobacter clade. Environmental Microbiology 17:5048–5062.

70. Ramamurthy T, Chowdhury G, Pazhani GP, Shinoda S. 2014. *Vibrio fluvialis*: an emerging human pathogen. Front Microbiol 5:91.

71. Weinstock MT, Hesek ED, Wilson CM, Gibson DG. 2016. *Vibrio natriegens* as a fast-growing host for molecular biology. Nature Methods 13:849–851.

72. Des Soye BJ, Davidson SR, Weinstock MT, Gibson DG, Jewett MC. 2018. Establishing a high-yielding cell-free protein synthesis platform derived from *Vibrio natriegens*. ACS Synth Biol 7:2245–2255.

73. Wiegand DJ, Lee HH, Ostrov N, Church GM. 2018. Establishing a cell-free *Vibrio natriegens* expression system. ACS Synth Biol 7:2475–2479.

74. Thoma F, Blombach B. 2021. Metabolic engineering of *Vibrio natriegens*. Essays Biochem 65:381–392.

75. Schada von Borzyskowski L. 2023. Taking synthetic biology to the Seas: From blue chassis organisms to marine aquaforming. ChemBioChem 24:e202200786.

76. Ellis GA, Tschirhart T, Spangler J, Walper SA, Medintz IL, Vora GJ. 2019. Exploiting the feedstock flexibility of the emergent synthetic biology chassis *Vibrio natriegens* for engineered natural product production. Mar Drugs 17.

77. Tschirhart T, Shukla V, Kelly EE, Schultzhaus Z, NewRingeisen E, Erickson JS, Wang Z, Garcia W, Curl E, Egbert RG, Yeung E, Vora GJ. 2019. Synthetic biology tools for the fast-growing Mmrine bacterium *Vibrio natriegens*. ACS Synth Biol 8:2069–2079.

78. VanArsdale E, Kelly E, Sayer CV, Vora GJ, Tschirhart T. 2024. Engineering xylose induction in *Vibrio natriegens* for biomanufacturing applications. Biotechnol Bioeng 121:3572–3581.

79. Fitzsimmons LF, Hampel KJ, Wargo MJ. 2012. Cellular choline and glycine betaine pools impact osmoprotection and phospholipase C production in *Pseudomonas aeruginosa*. J Bacteriol 194:4718–26.

80. Guzman LM, Belin D, Carson MJ, Beckwith J. 1995. Tight regulation, modulation, and high-level expression by vectors containing the arabinose PBAD promoter. J Bacteriol 177:4121–30.

81. Horton RM, Hunt HD, Ho SN, Pullen JK, Pease LR. 1989. Engineering hybrid genes without the use of restriction enzymes: gene splicing by overlap extension. . Gene 77:61–68.

82. Philippe N, Alcaraz JP, Coursange E, Geiselmann J, Schneider D. 2004. Improvement of pCVD442, a suicide plasmid for gene allele exchange in bacteria. Plasmid 51:246–55.

83. Dehio C, Meyer M. 1997. Maintenance of broad-host-range incompatibility group P and group Q plasmids and transposition of Tn5 in **Bartonella henselae** following conjugal plasmid transfer from *Escherichia coli*. J Bacteriol 179:538–40.

84. Thompson JD, Higgins DG, Gibson TJ. 1994. CLUSTAL W: improving the sensitivity of progressive multiple sequence alignment through sequence weighting, position-specific gap penalties and weight matrix choice. Nucleic Acids Res 22:4673–80.

85. Tamura K, Stecher G, Kumar S. 2021. MEGA11: Molecular Evolutionary Genetics Analysis Version 11. Mol Biol Evol 38:3022–3027.

